# Can one trust kinetic and thermodynamic observables from biased metadynamics simulations: detailed quantitative benchmarks on millimolar drug fragment dissociation

**DOI:** 10.1101/558601

**Authors:** Debabrata Pramanik, Zachary Smith, Adam Kells, Pratyush Tiwary

## Abstract

Obtaining atomistic resolution of ligand dissociation from a protein is a much sought after experimental and computational challenge. Structural details of the dissociation process are in general hard to capture in experiments, while the relevant timescales are far beyond molecular dynamics (MD) simulations even with the most powerful super-computers. As such many different specialized enhanced sampling methods have been proposed that make it possible to efficiently calculate the dissociation mechanisms in protein-ligand systems. However, accurate benchmarks against long unbiased MD simulations are either not reported yet or simply not feasible due to the extremely long timescales. In this manuscript, we consider one such recent method “infrequent metadynamics”, and benchmark in detail the various thermodynamics and kinetic information obtained from this method against extensive unbiased MD simulations for the dissociation dynamics of two different millimolar fragments from the protein FKBP in explicit water with residence times in nanoseconds to microseconds regime. We find that the metadynamics approach gives the same binding free energy profile, dissociation pathway and ligand residence time as the unbiased MD, albeit using only 6 to 50 times lower computational resources. Furthermore, we demonstrate how the metadynamics approach can self-consistently be used to ascertain whether the reweighted kinetic constants are reliable or not. We thus conclude that the answer to the question posed in the title of this manuscript is: statistically speaking, yes.

## I. INTRODUCTION

Protein-ligand interactions are some of the most common, yet diverse, processes in the human body, with direct relevance to our fundamental understanding of biology, as well as to the design of more effective and selective drugs for treating various diseases.^1–4^ For rational drug design, it becomes very desirable to understand the pathways through which ligands associate/dissociate, and the mechanisms at play in stabilizing and destabilizing these interactions.^5–8^ All-atom computer simulations have played a significant role in this avenue,^3,5,9–22^ but their true potential is arguably still limited due to the complexity of the problem.^22–25^ The complexity is at least two fold – (a) accuracy of classical force-fields, and (b) the rare event nature of the problem. In this work, we focus on the rare event nature of the problem, wherein straightforward classical molecular dynamics (MD) falls far short of the timescales needed to observe spontaneous dissociation of a ligand from the protein. Specialized computing hardware has also been used for reaching timescales in unbiased MD that would be unfathomable a few years ago.^26^ However, such approaches are computationally expensive, and difficult to deploy on a cost-effective basis for screening through multiple protein-ligand systems in a drug discovery program. As such, in order to observe protein-ligand dissociation, even with limited computational resources, numerous enhanced sampling methods have been proposed over the years.^27,28^

These sampling methods have been proposed both for calculating binding affinities^29–32^ and association/dissociation rate constants.^23,33–37^ Their reliability is less arguable in the context of binding affinity calculations (especially relative binding affinities), even though there is still scope for improvement. What remains an open question is whether such sampling methods can be trusted, while still being computationally advantageous over unbiased methods, if the objective is to calculate dissociation mechanisms and associated timescales. Thus the reliability of such sampling methods remains not entirely settled, primarily due to the lack of accurate benchmarking against extremely long unbiased MD simulations. Any benchmarks reported so far are against experimental measurements of the dissociation rate constant. While these are encouraging, they could very well be so due to cancellation of errors.

In this work we consider the unbinding of two different fragments from the protein FKBP. We perform extensive long (close to one hundred microseconds in total) unbiased MD simulations in-house to obtain accurate statistics of the timescales and the pathways. In parallel, we perform infrequent metadynamics^33^ to get these observables, which is between 6 to 50 times faster on average for the two ligands. These simulations are performed using the exact same force-fields for protein, ligand and water, and thermostat/barostat parameters. A key ingredient of infrequent metadynamics is knowing a fairly accurate reaction coordinate (RC) which is used to perform the biasing. Here we find such a RC in an automated manner through the use of the RC optimization method Spectral gap optimization of order parameters (SGOOP).^34–36^ Through the combination of these two methods we demonstrate that we can calculate dissociation pathways and timescales well within error margin agreement with our unbiased MD estimates, albeit in two orders of magnitude lesser computer time. Furthermore we demonstrate when the combined approach of SGOOP and infrequent meta-dynamics is reliable, when it isn’t, and how we can successfully tell the difference between both the cases even if we don’t know the true answer.^38^

We believe this is one of the first such reported works that carefully benchmarks and demonstrates the utility of infrequent metadynamics-based approach for calculates ligand dissociation mechanisms. Given the rapidly increasing popularity of this approach, this work serves an important purpose in demonstrating that this approach can indeed be used reliably.

## II. THEORY

### A. Spectral gap optimization of order parameters (SGOOP)

This is a recently proposed method to construct an optimized RC as a linear or non-linear combination of a pre-specified dictionary of order parameters,^34–36^ which can then be used to perform enhanced sampling. It has been shown to be especially useful for rare event systems.^22,35,36^ To use SGOOP, the key inputs are (a) stationary probability density estimate of the system obtained either through a very long unbiased MD run when feasible, or by performing sampling along a putative RC, (b) suitable dynamical observables or constraints on the system, which could for instance be the average number of transitions in a unit time. We refer the reader to Ref. 35 as well as the Supplemental Information (SI) for specific details of these constraints. Given these two pieces of information, SGOOP uses a maximum path entropy (or Caliber) model^39–41^ to calculate the transition probability matrix along any given trial RC. The spectral gap of this transition probability matrix^34,35^ serves as a measure of the quality of the trial RC. The central motivation is that the optimal RC (i) should be able to separate out as many metastable states as possible, (ii) any degrees of freedom in the full system not encapsulated by this RC in terms of discernible barriers should be relatively as fast as possible. By then using simulated annealing to optimize the spectral gap as a function of the weights of different order parameters contributing to the RC, SGOOP then learns the optimal, RC given the stationary and dynamic data at hand. This RC can then be used to perform efficient and reliable enhanced sampling, for instance infrequent metadynamics^33^ which we describe next.

### B. Infrequent metadynamics

In its traditional form, the central idea behind metadynamics^42^ is that by adding a time-dependent bias to a given system, it can be encouraged to gradually escape free energy minima that an unbiased simulation would get trapped in, and instead explore various facets of the energy landscape. Through a reweighting^43^ procedure one can then construct the true underlying Boltzmann probability distribution as a function of any order parameter. In its more recent so-called “infrequent” version, it is possible to also calculate unbiased kinetic rate constants and dissociation pathways from biased simulations.^33^ The idea here is to make the bias deposition infrequent relative to the time spent in the short-lived transition state when crossing between any two metastable states. This way it can be shown that (a) metadynamics preserves the unbiased state-to-state sequence of transitions, (b) the unbiased timescales for the state-to-state escape time can be recovered through the calculation of an acceleration factor (SI). Furthermore, the reliability of these reweighted timescales can be verified a posteriori through a Kolmogorov-Smirnov test,^38^ wherein a p-value higher than 0.05 indicates reliability (SI).

## III. RESULTS

The central idea in this work is to verify the reliability of a combination of the above two methods on a pair of specific ligands with millimolar affinity, dissociating from the FKBP protein. These ligands have dissociation timescales in the regime that very accurate benchmarks can be obtained through unbiased MD.^44–46^ Yet, these are slow enough that using infrequent metadynamics on SGOOP-optimized RC one can achieve meaningful computational speed-up of up to 50 times. The question we then ask is: how reliable are the kinetic and thermodynamic observables obtained from metadynamics, relative to the very well converged benchmarks from in-house unbiased MD using identical force-field, thermostat, barostat etc. These observables, which we now cover in a sequential manner, include the (i) residence time, (ii) dissociation pathway and (iii) binding free energy profile. We first describe the protocol followed here. Further details about the systems, their preparation and the MD simulation parameters themselves are provided in the SI. We would like to point out that our initial attempt was to reproduce the unbiased estimates reported in Ref. 47. However, we found that small differences in ligand and other parameterizations can lead to profound differences in the kinetics. As such, here we report our own in-house unbiased MD for benchmark. It is worth pointing out that the relative trend between the two ligands through our unbiased MD is the same as that reported in Ref. 47.

### A. Protocol

For each system, we first constructed a dictionary of order parameters, comprising 10 different protein-ligand distances (Table I). These are fairly generic order parameter choices and their selection represents the only non-automated part of this work. For the stationary probability density input into SGOOP,^34^ we are presented with 2 choices – either (i) from metadynamics along a trial RC followed by reweighting, or (ii) a long unbiased MD run. Here we report results using (i) for the slower ligand and (ii) for the faster ligand, but do not expect this to be crucial especially from the perspective of infrequent metadynamics. This trajectory serves as both the stationary and dynamical input to SGOOP (SI), with which we obtained an optimized RC, detailed in Table I. With this SGOOP-optimized RC, we perform many independent infrequent metadynamics runs along this optimized RC to get dissociation of the ligand from the primary binding site. In any metadynamics simulation, an important control parameter is the bias deposition frequency which can be tuned to make the enhanced sampling more, or less aggressive. Here, we use a bias deposition frequency of once every 8 ps to calculate free energies, and two different bias deposition frequencies to perform infrequent metadynamics for calculating kinetics. These are once every 8 ps or once every 60 ps. Other metadynamics and MD parameters are given in detail in SI. To calculate the average binding free energy surface (FES) from many independent unbiased MD simulations, we calculate normalized probability for each run and do averaging over probability to get average FES. Similarly for metadynamics, we calculate the unbiased probability by using reweighting method^43^ for each run and then average over probability to get average FES.

**TABLE I:** SGOOP builds a low-dimensional RC starting from a pre-selected larger dictionary of order parameters. Here we provide the constituents of this dictionary for both systems, along with their initial and optimized weights. The two ends of the ligand, A and B have been shown in Fig. 1 in SI.

| Order<br>parameter | Type | Definition | Initial<br>weights<br>for BUT | Optimized<br>weights<br>for BUT | Initial<br>weights<br>for DAP | Optimized<br>weights<br>for DAP |
| --- | --- | --- | --- | --- | --- | --- |
| $d_1$ | protein-ligand<br>distance | Trp59-ligand | 1 | 1 | 1 | -1 |
| $d_2$ | protein-ligand<br>distance | Val55-ligand | 0 | 0.5761 | 0 | 0.6031 |
| $d_3$ | protein-ligand<br>distance | Phe46-ligand | 0 | 0.2047 | 0 | 0.5414 |
| $d_4$ | protein-ligand<br>distance | Tyr26-ligand | 0 | 0.2128 | 0 | -0.6091 |
| $d_5$ | protein-ligand<br>distance | Ile56-ligand | 0 | 0.1839 | 0 | -0.0121 |
| $d_6$ | protein-ligand<br>distance | Phe99-ligand | 0 | -0.3572 | 0 | 0.6075 |
| $d_7$ | protein-ligand<br>distance | Trp59-end A | 0 | 0.4083 | 0 | 0.5463 |
| $d_8$ | protein-ligand<br>distance | Trp59-end B | 0 | -0.6623 | 0 | 0.3385 |
| $d_9$ | relative protein-ligand distance | $d_7-d_8$ (relative<br>distance of Trp59<br>from the two ends<br>of the ligand) | 0 | 0.3245 | 0 | 0.4930 |
| $d_{10}$ | intra-ligand distance | ligand end A to<br>end B distance | 0 | 0.9379 | 0 | -0.2723 |

### B. Residence time

For reasons that will become apparent to the reader in Sec. III C, the residence time in this work is defined as the average time a ligand takes to reach the solvent exposed surface of the protein, starting from the bound pose. Here it is defined as *d*_1_ > 1.8 *Å* (Table I). Given the inherent roughness of the protein, this is an approximate *ad hoc* estimate. However, we emphasize that since we are comparing unbiased MD with metadynamics, the exact value of this parameter is not a cause for concern as long as we are consistent between both schemes. For unbiased MD, the residence time is measured by averaging over 20 independent runs starting from bound pose with randomized velocities. Similar procedure is followed for the metadynamics runs, but after taking the acceleration factor into account to correct for the effect of bias (SI).

For BUT (4-hydroxy-2-butanone), we obtain a residence time of 21.3±0.2 ns from unbiased MD, and 27.3±0.1 ns from infrequent metadynamics, which demonstrates excellent agreement especially taking into account that the metadynamics runs had acceleration factors ranging from 1.3 to 11.2, averaging around 6. The p-value for the metadynamics derived timescales when fitted using the KS test from Ref. 38(SI) was 0.82, far above the cut-off of 0.05. This further demonstrates the reliability of the metadynamics derived timescales even if one did not have access to unbiased MD values. For DAP (4-diethylamino-2-butanone), we obtain a residence time of 0.54 *µ*s from unbiased MD, and 1.83±0.03 *µ*s from infrequent metadynamics (see SI for further details of the respective statistics and further statistical analyses). Here we need to take into account that (a) the metadynamics runs had an average acceleration factor of 51 over the unbiased runs, varying between 1.2 to 375.9 and (b) seven of the twenty unbiased runs went to as much as 2 *µ*s without dissociating, due to which we do not assign error bars to the unbiased estimate. In light of the high computational speed-up, and considering that the unbiased estimate is strictly a lower bound due to undissociated runs, this again demonstrates excellent agreement between both approaches. The averaged p-value for the metadynamics derived timescales when fitted using the KS test from Ref. 38 was 0.08 (SI), again above the cut-off of 0.05. Interestingly, the use of biasing every 8 ps was sufficient for the BUT system to obtain p-value above 0.05. The use of the same biasing frequency as well as gentler frequency of once every 20 ps gave a p-value much smaller than 0.05 for DAP, and thus this system was biased more infrequently at once every 60 ps where we obtained p-value of 0.08, which is above the cut-off of 0.05. It is interesting that even though the p-value is only slightly above the cut-off, and poorer than that for BUT, the agreement between unbiased and metadynamics timescale estimates is quite good considering the high value of the acceleration factor and that 35 % of unbiased runs did not even dissociate in 2 *µ*s each.

### C. Dissociation pathways

While the residence time is a very important metric for ligand dissociation, an equally if not even more important metric is the dissociation pathway adopted by the ligand. Thus, here we ask the question concerning the differences and similarities in the dissociation pathways for both ligands as seen through infrequent metadynamics and through the benchmark long unbiased MD simulations. As can be seen from Fig. 1, the results are reassuring and excellent. In this figure we have provided representative superimposed snapshots from typical dissociation trajectory for each ligand, calculated through unbiased MD and through infrequent metadynamics. For both simulation methods, we observe that the ligand dissociation here is a combination of two sequential steps. First, the ligand takes a well-defined path from the bound pose to the solvent exposed surface of the protein. A useful image is that of a snorkeler coming to the surface of the ocean. Then, the remainder of the dissociation is entropically driven wherein the ligand can take various different pathways on its way to being fully solvated.

**FIG. 1:**
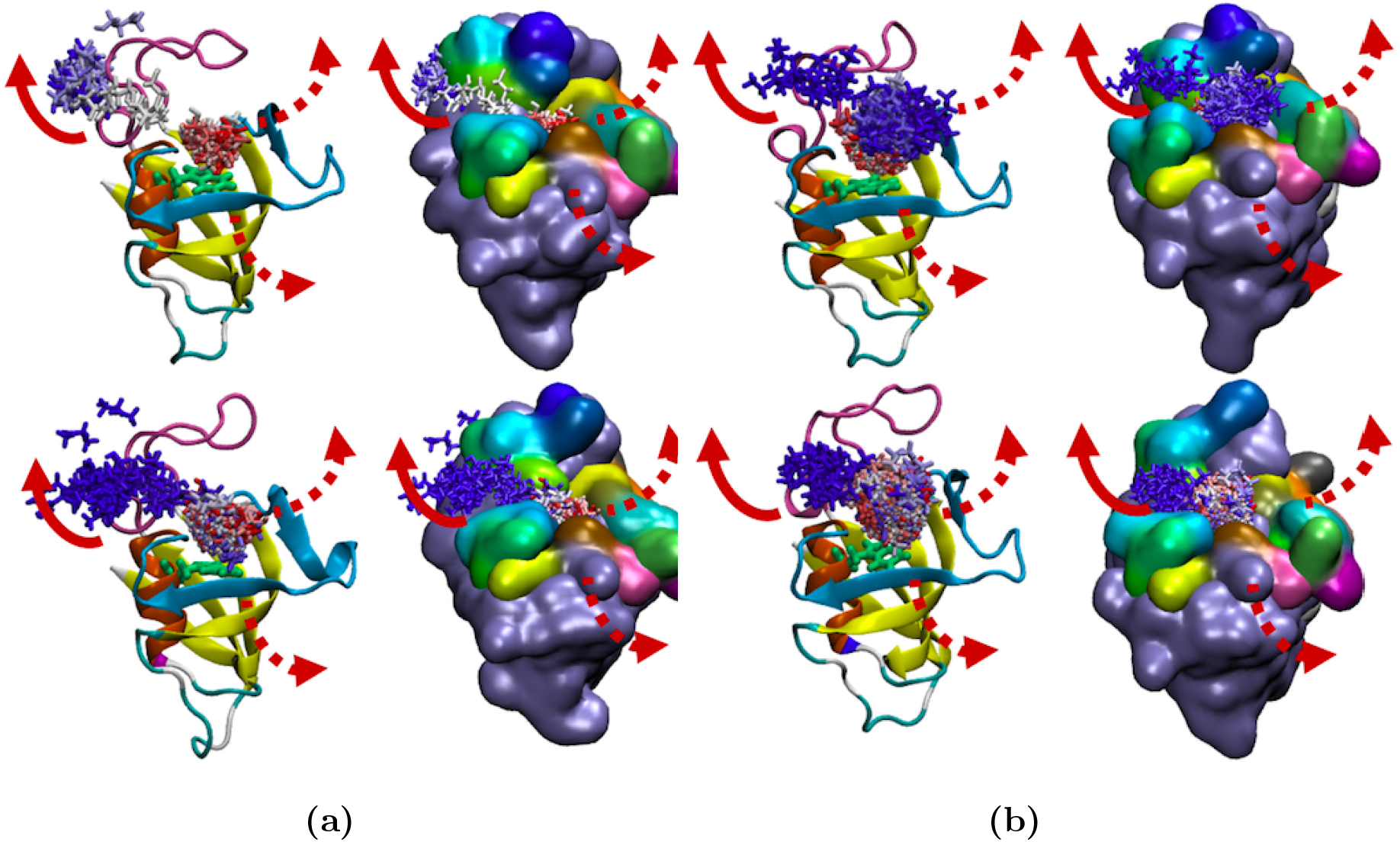
Here (a) and (b) are for the ligands BUT (4-hydroxy-2-butanone) and DAP (4-diethylamino-2-butanone) respectively dissociating from the primary binding site of the protein FKBP. As described in Sec. III C, the dissociation comprises two sequential events, first a slow part wherein the ligand comes to the solvent-exposed surface of the protein, and a second relatively much faster part, when it diffuses on the surface before finally reaching the solvent. In (a) as well as (b), one representative pathway has been shown here in cartoon (left) and surface (right) representations by solid arrows. Other observed pathways, differing only in the second surface diffusion part, are shown by dashed arrows. Top and bottom rows show results from unbiased MD and infrequent metadynamics respectively. For all cases, important segments of the protein near the binding site have been highlighted by different colors. Location of the binding pocket has been shown by the residue Trp which is in the binding pocket highlighted by green color (licorice representation).

Our central finding is that metadynamics and unbiased MD preserve the first step in dissociation, which is the pathway taken to come to the surface. After that there are entropically many pathways and there is no clean statistics on a single preferred final diffusion mechanism, as one would expect. This is clearly illustrated in Fig. 1. Here we show the dissociation pathways for both the ligands, namely BUT (Fig. 1(a)) and DAP (Fig. 1(b)). The top row is from unbiased MD and the bottom row from infrequent metadynamics. In both (a) and (b), the left sub-figure shows a secondary structure view of the protein, highlighting for reference the residue Trp59. The right sub-figure shows the exit point on the surface protein where the ligand first appears, followed by a typical dissociation trajectory after this point. As can be seen here, for both ligands the pathway until the first appearance of the the ligand on the surface of the protein is preserved between infrequent metadynamics and unbiased MD. Furthermore, the region on the surface of the protein where either ligand first makes an appearance is also preserved for both methods. This is the more crucial part of the dynamics and it is a clear evidence for the potential of infrequent metadynamics in recovering dissoication pathways. As for the pathway after exit to surface, we believe it will take on the order of 100, perhaps more, dissociation trajectories to accurately compare their statistics. Thus here we observe this second and relatively much faster part of the dissociation only qualitatively. In Fig. 1(a) we show one such representative direction the ligand takes from the surface top to the aqueous medium. Similarly, in Fig. 1(b) we show representative pathway for ligand DAP from the bound pose to the surface of the protein and then from the surface to the aqueous medium.

### D. Free energy profile

Last, but not the least, we compare the two methods by analyzing the binding free energy profiles obtained using them. In Fig. 2 we provide these free energy profiles as a function of the SGOOP-optimized RC for both ligands, calculated through unbiased MD and through metadynamics for different frequencies. The agreement between metadynamics and unbiased MD is again well within error bars. Furthermore, as can be expected, making metadynamics more infrequent leads to improved agreement for both the ligands.

**FIG. 2:**
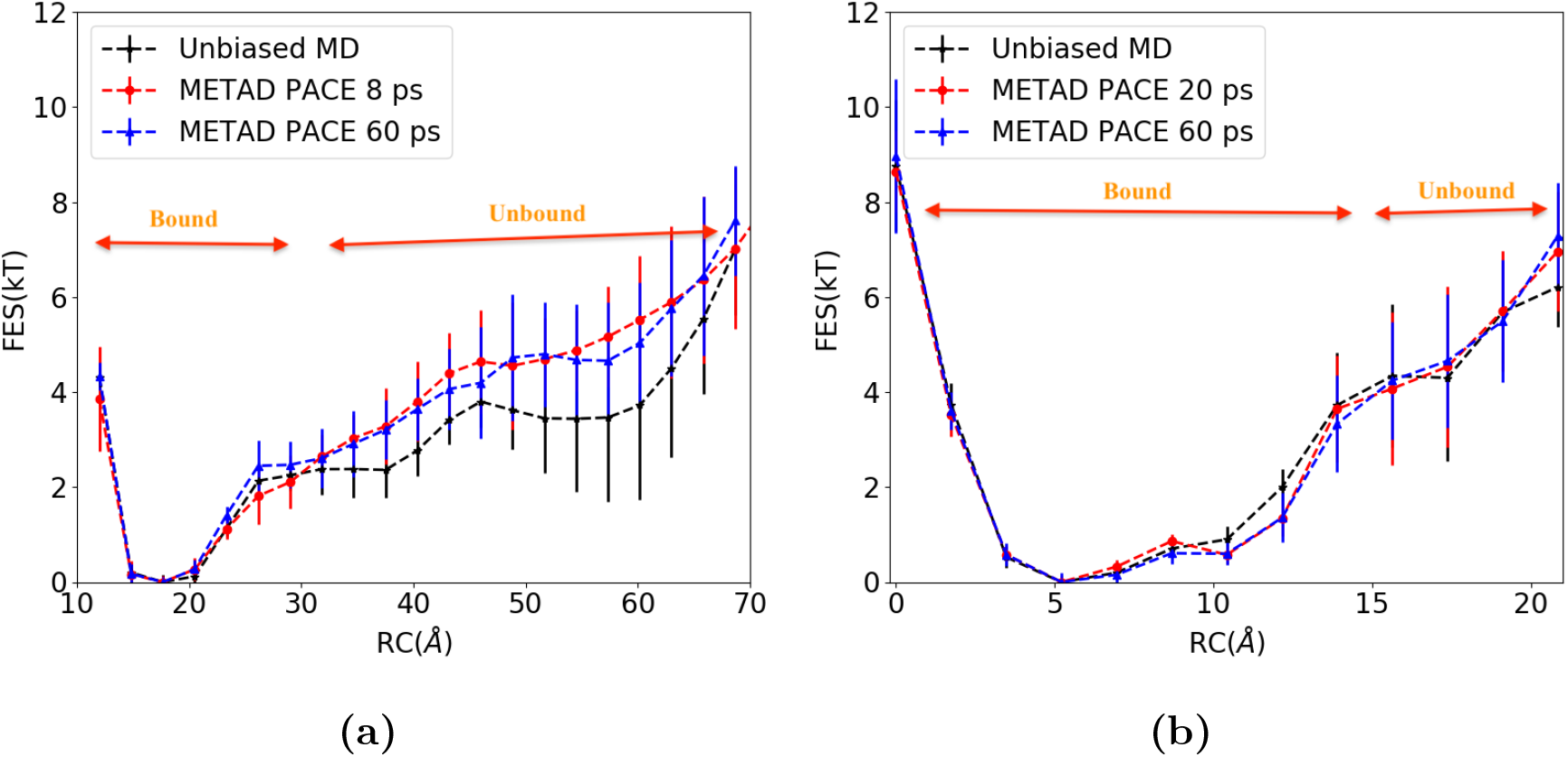
(a) Averaged free energy surface (FES) for BUT from unbiased MD (black) and metadynamics (other colors) with corresponding error bars from independent runs. We show two FES for metadynamics with different bias depositing pace (or inverse of biasing frequency), with fast and slow biasing paces denoted by red and blue colors respectively. All free energies have been averaged over sufficiently large number of independent data to have more accurate statistics, equaling 28 and 21 respectively for unbiased MD and metadynamics respectively. The biasing paces are 8 ps (red), and 60 ps (blue). (b) Averaged FES for DAP from unbiased MD (black) (20 independent runs), and metadynamics. For metadynamics we had 36 independent runs with pace 20 ps (red), and 25 with pace 60 ps (blue).

## IV. DISCUSSION

Protein-ligand dissociation is a problem of great significance in chemistry and biology, for which a number of theoretical^48^ as well as specialized sampling methods^49^ have been proposed that surmount the timescale limitations of unbiased all-atom MD. An open question in the field has been to ascertain the veracity of such methods by benchmarking against detailed unbiased MD calculations. Towards this end, in this work we have studied the dissociation of two ligands BUT and DAP from the primary binding site of the protein FKBP, using extensive unbiased MD calculations nearing a total of around 100 *µ*s, as well as much faster infrequent metadynamics-based calculations. For both methods, we obtained details of dissociation pathway, residence time and free energy profile, while using identical force field and other simulations parameters for both the schemes. A central requirement for the infrequent metadynamics scheme is having an approximate reaction coordinate along which to the bias the system. Here we found such a reaction coordinate in a fairly automated manner using the SGOOP method proposed by Tiwary and Berne,^34^ as a function of a larger dictionary of 10 order parameters. Our key finding is that the SGOOP and infrequent metadynamics-based approach can give fairly accurate estimates of various kinetic and thermodynamic observables with much less computational effort than a brute-force MD approach. For the slower ligand DAP with residence time around 2 *µ* s, the maximal computational speed-up was 50 times when using a biasing frequency of once every 60 ps. Attempting to bias more aggressively led to higher speed-ups but inaccurate kinetics. For the faster ligand BUT with residence time around 27 ns, the computational speed-up was 6 times relative to unbiased MD. Another interesting finding is that using the approach of Salvalaglio et al.^38^ we could self-consistently, and without knowing the true residence time, determine which biasing frequency was too aggressive or not. The infrequent metadynamics-based residence times for DAP and BUT are in excellent agreement with estimates from unbiased MD, as well as in qualitative agreement with the numbers reported by Pan et al using different force-field parametrizations.^47^ Finally, we demonstrated that the potential of such a metadynamics-based approach goes beyond residence time, but also includes accurate characterization of the dissociation pathway as well as the dissociation free energy profile. We thus conclude that there is promise for the use of recent enhanced sampling-based approaches such as SGOOP, infrequent metadynamics and others^49^ in studying ligand dissociation mecnahisms in all-atom resolution, but at significantly lower computational expense than unbiased molecular dynamics.

## ACKNOWLEDGMENTS

We thank Deepthought2, MARCC and XSEDE (projects CHE180007P and CHE180027P) for providing the computational resources used to perform this work. PT would like to thank University of Maryland Graduate School for financial support through the Research and Scholarship Award (RASA). We thank Albert Pan for providing the starting structures and parameters for the systems, and Michael Shirts for help with converting DESMOND files to GROMACS. A. K. is supported by the EPSRC Centre for Doctoral Training in Cross-Disciplinary Approaches to Non-Equilibrium Systems (CANES, EP/ L015854/1)

## REFERENCES

1. P. J. Hajduk and J. Greer, Nature Reviews Drug Discovery 6, 211 (2007).

2. M. Congreve, G. Chessari, D. Tisi, and A. J. Woodhead, Journal of Medicinal Chemistry 51, 3661 (2008).

3. R. O. Dror, R. M. Dirks, J. Grossman, H. Xu, and D. E. Shaw, Annual Review of Biophysics 41, 429 (2012).

4. J. G. Zeevaart, L. Wang, V. V. Thakur, C. S. Leung, J. Tirado-Rives, C. M. Bailey, R. A. Domaoal, K. S. Anderson, and W. L. Jorgensen, Journal of the American Chemical Society 130, 9492 (2008).

5. R. O. Dror, A. C. Pan, D. H. Arlow, D. W. Borhani, P. Maragakis, Y. Shan, H. Xu, and D. E. Shaw, Proceedings of the National Academy of Sciences 108, 13118 (2011).

6. K. J. Kohlhoff, D. Shukla, M. Lawrenz, G. R. Bowman, D. E. Konerding, D. Belov, R. B. Altman, and V. S. Pande, Nature Chemistry 6, 15 (2014).

7. J. Mortier, C. Rakers, M. Bermudez, M. S. Murgueitio, S. Riniker, and G. Wolber, Drug Discovery Today 20, 686 (2015).

8. D. Sabbadin and S. Moro, J. Chem. Inf. Model. 54, 372 (2014).

9. C. Simmerling, B. Strockbine, and A. E. Roitberg, Journal of the American Chemical Society 124, 11258 (2002).

10. S. Y. Noskov, S. Berneche, and B. Roux, Nature 431, 830 (2004).

11. M. M. Seibert, A. Patriksson, B. Hess, and D. Van Der Spoel, Journal of molecular biology 354, 173 (2005).

12. J. D. Faraldo-Gómez and B. Roux, Proceedings of the National Academy of Sciences 104, 13643 (2007).

13. A. Berteotti, A. Cavalli, D. Branduardi, F. L. Gervasio, M. Recanatini, and M. Parrinello, Journal of the American Chemical Society 131, 244 (2008).

14. R. O. Dror, D. H. Arlow, D. W. Borhani, M. Ø. Jensen, S. Piana, and D. E. Shaw, Proceedings of the National Academy of Sciences 106, 4689 (2009).

15. S. Vanni, M. Neri, I. Tavernelli, and U. Rothlisberger, Biochemistry 48, 4789 (2009).

16. E. Lyman, C. Higgs, B. Kim, D. Lupyan, J. C. Shelley, R. Farid, and G. A. Voth, Structure 17, 1660 (2009).

17. V. A. Voelz, G. R. Bowman, K. Beauchamp, and V. S. Pande, Journal of the American Chemical Society 132, 1526 (2010).

18. G. Enkavi and E. Tajkhorshid, Biochemistry 49, 1105 (2010).

19. D. E. Shaw, P. Maragakis, K. Lindorff-Larsen, S. Piana, R. O. Dror, M. P. Eastwood, J. A. Bank, J. M. Jumper, J. K. Salmon, Y. Shan, et al., Science 330, 341 (2010).

20. J. Mondal, N. Ahalawat, S. Pandit, L. E. Kay, and P. Vallurupalli, PLoS computational biology 14, e1006180 (2018).

21. J. M. L. Ribeiro, S.-T. Tsai, D. Pramanik, Y. Wang, and P. Tiwary, Biochemistry 58, 156 (2019).

22. P. Tiwary, J. Phys. Chem. B 121, 10841 (2017).

23. P. Tiwary, V. Limongelli, M. Salvalaglio, and M. Parrinello, Proceedings of the National Academy of Sciences 112, E386 (2015).

24. R. A. Copeland, Nature Reviews Drug Discovery 15, 87 (2016).

25. Y. Shan, E. T. Kim, M. P. Eastwood, R. O. Dror, M. A. Seeliger, and D. E. Shaw, J. Amer. Chem. Soc. 133, 9181 (2011), http://dx.doi.org/10.1021/ja202726y.

26. D. E. Shaw, M. M. Deneroff, R. O. Dror, J. S. Kuskin, R. H. Larson, J. K. Salmon, C. Young, B. Batson, K. J. Bowers, and J. C. Chao, Communications of the ACM 51, 91 (2008).

27. P. Tiwary and A. van de Walle, in Multiscale Materials Modeling for Nanomechanics (Springer, 2016) pp. 195–221.

28. O. Valsson, P. Tiwary, and M. Parrinello, Ann. Rev. Phys. Chem. 67, 159 (2016).

29. Y. Deng and B. Roux, J. Phys. Chem. B 113, 2234 (2009).

30. L. Wang, Y. Wu, Y. Deng, B. Kim, L. Pierce, G. Krilov, D. Lupyan, S. Robinson, M. K. Dahlgren, J. Greenwood, et al., Journal of the American Chemical Society 137, 2695 (2015).

31. L. Wang, B. J. Berne, and R. A. Friesner, Proceedings of the National Academy of Sciences 109, 1937 (2012).

32. V. Limongelli, M. Bonomi, and M. Parrinello, Proceedings of the National Academy of Sciences (2013), 10.1073/pnas.1303186110.

33. P. Tiwary and M. Parrinello, Phys. Rev. Lett. 111, 230602 (2013).

34. P. Tiwary and B. J. Berne, Proc. Natl. Acad. Sci. 113, 2839 (2016).

35. Z. Smith, D. Pramanik, S.-T. Tsai, and P. Tiwary, J. Chem. Phys. (2018), 10.1063/1.5064856.

36. P. Tiwary and B. J. Berne, J. Chem. Phys. 145, 054113 (2016).

37. L. Mollica, S. Decherchi, S. R. Zia, R. Gaspari, A. Cavalli, and W. Rocchia, Scientific Reports 5, 11539 (2015).

38. M. Salvalaglio, P. Tiwary, and M. Parrinello, J. Chem. Theor. Comp. 10, 1420 (2014).

39. E. T. Jaynes, Ann. Rev. Phys. Chem. 31, 579 (1980).

40. S. Presśe, K. Ghosh, J. Lee, and K. A. Dill, Rev. Mod. Phys. 85, 1115 (2013).

41. P. D. Dixit, A. Jain, G. Stock, and K. A. Dill, J. Chem. Theor. Comp. 11, 5464 (2015).

42. A. Laio and M. Parrinello, Proc Natl Acad Sci 99, 12562 (2002).

43. P. Tiwary and M. Parrinello, J. Phys. Chem. B 119, 736 (2014).

44. D. Huang and A. Caflisch, PLoS Comput. Biol. 7, e1002002 (2011).

45. D. Huang and A. Caflisch, ChemMedChem 6, 1578 (2011).

46. A. C. Pan, H. Xu, T. Palpant, and D. E. Shaw, Journal of Chemical Theory and Computation 13, 3372 (2017).

47. A. C. Pan, H. Xu, T. Palpant, and D. E. Shaw, J. Chem. Theor. Comp. 13, 3372 (2017).

48. N. Agmon and J. Hopfield, J. Chem. Phys. 79, 2042 (1983).

49. J. M. L. Ribeiro, S.-T. Tsai, D. Pramanik, Y. Wang, and P. Tiwary, Biochemistry 58, 156 (2018).

