## Supplement for "Can one trust kinetic and thermodynamic observables from biased metadynamics simulations: detailed quantitative benchmarks on millimolar drug fragment dissociation"

(Dated: February 25, 2019)

In this supplement, we provide further information on the simulations in an itemized manner.

1. **System parameterization and simulation set-up**

   (a) Starting structures for simulations were generated from files provided by Albert Pan, using in recent work[1]. These structures were converted from Desmond to Gromacs format using the software InterMol (github.com/mrshirts/InterMol). Below in Fig. 1(a)-(b) respectivly, we show the chemical compositions of the two ligands, BUT and DAP.

   (b) For the protein FKBP, the Amber99SB-ILDN force field was used with TIP3P water. The ligand was parameterized with the generalized amber force field (GAFF).

   (c) Equilibration was performed in the NPT ensemble for 1 ns. Production runs are then performed with Nose-Hoover thermostat at room temperature (300 K) and with a 2 fs timestep.

   (d) Well-tempered metadynamics and MD parameters: In metadynamics simulation for BUT, the biases were deposited with an initial hill height of 1.5 kJ, width 0.05 $\AA$, biasfactor 10 , and for DAP metadynamics the biases were deposited with a hill-height 0.75 kJ, width 0.05 $\AA$, biasfactor 5, at temperature 300 K. The biasing frequencies varied and have already been detailed in the main text.

2. **Suitable dynamical observable or constraint on the system, which could for instance be the average number of transitions in unit time:** In SGOOP, to calculate the transition rate matrix, we need to choose a dynamical observable which can be used in the Maximum Caliber framework. Below in Eq.1, we show an element of the transition rate matrix. To get it, first we take a short unbiased MD trajectory for the set of order parameters and then we discretize the putative RC in n integral intervals. If $\langle N \rangle$ is the average number

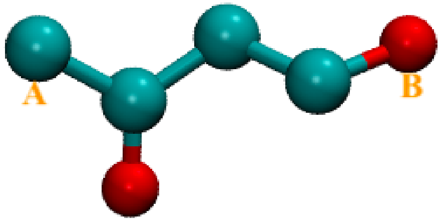


**(a)**

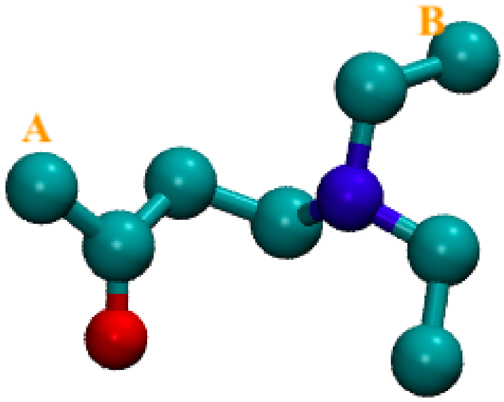


**(b)**

**FIG. 1:** Chemical composition of (a) BUT (4-hydroxy-2-butanone) and (b) DAP (4-diethylamino-2-butanone). The heavy atoms (carbon (green), oxygen (red), nitrogen (blue)) have been shown without hydrogen atoms for both ligands.

of nearest-neighbour transitions along the putative RC intervals in unit time $\Delta t$, we use this as the dynamical observable in the rate matrix calculation. Here, the rate of going from grid site m to n is given by Eq.1, where $p_m$ and $p_n$ are the unbiased stationary probabilities for the systems of being at grid site m and n. Thus we can get the full transition rate matrix and calculate all the eigenvalues of this matrix. The eigenvalues can be use to calculate the spectral gap.

$$k_{mn} = \left\{ \frac{\langle N \rangle}{\sum \sqrt{p_n p_m}} \right\} \sqrt{\frac{p_n}{p_m}} \tag{1}$$

3. **SGOOP derived eigenspectrums and top few eigenvectors for both systems** For BUT SGOOP obtains a self-consistent 2-well solution with 1 barrier, and for DAP 3-well solution with 2 barriers. Here, the threshold value has been used as 1 kT, which means when the barrier height > 1 kT, we consider the newly indentified local minimum as a new metastable state. In Fig. 2(a) we mark the spectral gap with red arrows for BUT, and in 2(b) we show the free energy profile with stationary, $1^{st}$ and $2^{nd}$ eigenvectors. In Fig.3(a) we mark the spectral gap with red arrows for DAP, and in 3(b) we show the free energy profile with stationary, $1^{st}$, $2^{nd}$ and $3^{rd}$ eigenvectors. It can be seen that for BUT and DAP respectively, the $2^{nd}$ and $3^{rd}$ eigenvectors correspond to fast, intra-basin movements.

4. **Reweighting timescales from metadynamics through an acceleration factor approach:** The accelerated time corresponding to a metadynamics run is defined as $t_{acc}$=$\int_0^t \exp(\beta V(t))\delta t$. Here, $\beta$=1/$k_B$T, where $T$ is simulation temperature, $k_B$ is the Boltzmann constant, $V(t)$ is deposited bias as function of time and $\delta t$ is the unbiased integration time-step or a multiple thereof.

5. **Statistical reliability of the dynamics reweighted from metadynamics:** Here we provide detailed results of the Kolmogorov-Smirnov test from Ref. [2]. The key idea here is to examine the distribution of transition times for a given rare event obtained through reweighting[3] from the time-dependent biasing protocol of metadynamics. If the metadynamics biasing was infrequent enough to leave the transition states bias free, this should manifest in the transition time distribution being fit to a memoryless Poisson distribution. This in turn can bequantified through a Kolmogorov-Smirnov test, wherein a p-value higher than 0.05 indicates reliability of the dynamics reweighted from metadynamics.

   In order to make sure that there is no dependence we sample size, we selected at random 20 sets of 20 transition times with replacement from the infrequent metadynamics runs for BUT as well as for DAP. For DAP we had a total of 25 dissociation events through infrequent metadynamics while for BUT this number was 21. The choice of studying 20 transition times came from the observation that we had a total of 20 unbiased dissociation runs for the slower unbinder, namely DAP.

   As we mentioned in the main text, for BUT we were able to obtain a p-value above the cut-off of 0.05 even with an aggressive biasing frequency of once every 8 ps. Fig. shows the distribution of poisson fits over the 20 sets of randomly sampled 20 transition events (thus leaving 1 out in each set, as we had a total of 21 dissociation events). The average p-value was 0.82. The computational speed-up for this system averaged 6, while the transition times are within 30% of each other.

   For the second ligand DAP we were unable to obtain a p-value above the cut-off of 0.05 with the aggressive biasing frequency of once every 8 ps as well as once every 20 ps. Finally with biasing once every 60 ps we got a p-value above the cut-off. Fig. shows the distribution of poisson fits over the 20 sets of randomly sampled 20 transition events (thus leaving 5 out in each set, as we had a total of 25 dissociation events). The average p-value was 0.08. The computational speed-up for this system averaged 51, while the transition times are within 4 times of each other, which is quite satisfactory given that 7 out of 20 unbiased runs reached nearly 2 $\mu$s without dissociating.

---

[1] A. C. Pan, H. Xu, T. Palpant, and D. E. Shaw, Journal of Chemical Theory and Computation **13**, 3372 (2017).
[2] M. Salvalaglio, P. Tiwary, and M. Parrinello, J. Chem. Theor. Comp. **10**, 1420 (2014).
[3] P. Tiwary and M. Parrinello, Phys. Rev. Lett. **111**, 230602 (2013).

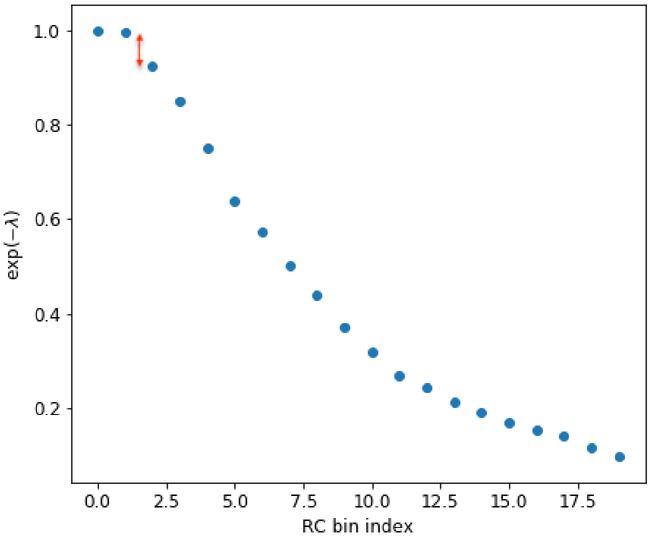


**(a)**

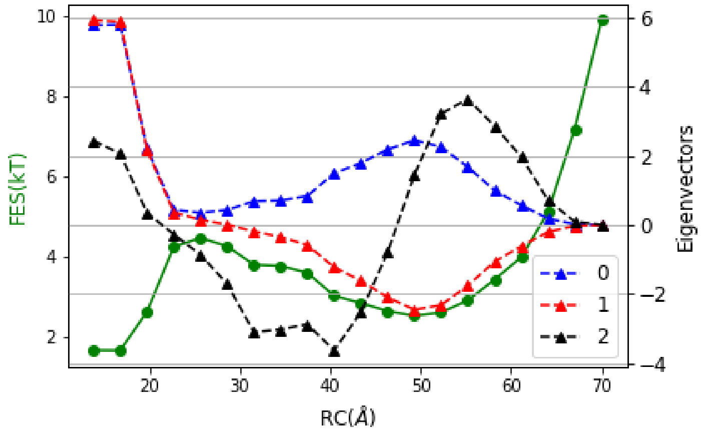


**(b)**

**FIG. 2:** (a) Plot of $\exp(-\lambda)$ with RC bin index for BUT. Here eigenvalues of Eq. 1 are denoted by $\lambda$. Red arrow shows the spectral gap in the plot. (b) FES (green color), stationary state (denoted by '0'), $1^{st}$ ('1') and $2^{nd}$ ('2') eigenvectors are shown.

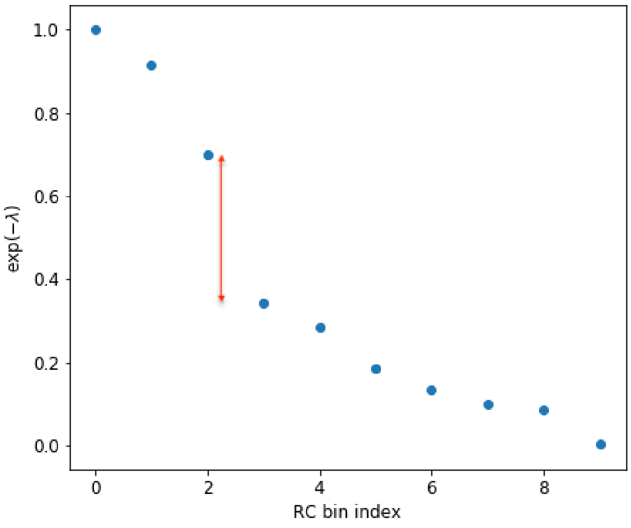


**(a)**

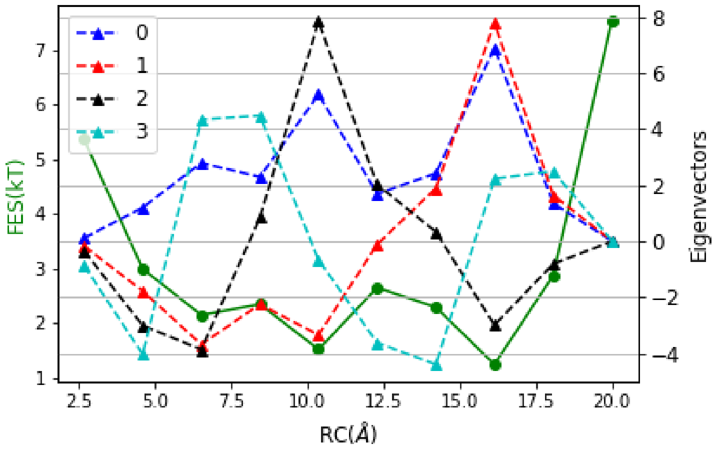


**(b)**

**FIG. 3:** (a) Plot of $\exp(-\lambda)$ with RC bin index for DAP. Here eigenvalues of Eq. 1 are denoted by $\lambda$. Red arrow shows the spectral gap in the plot. (b) FES (green color), stationary state (denoted by '0'), $1^{st}$ ('1'), $2^{nd}$ ('2') and $3^{rd}$ ('3') eigenvectors are shown.

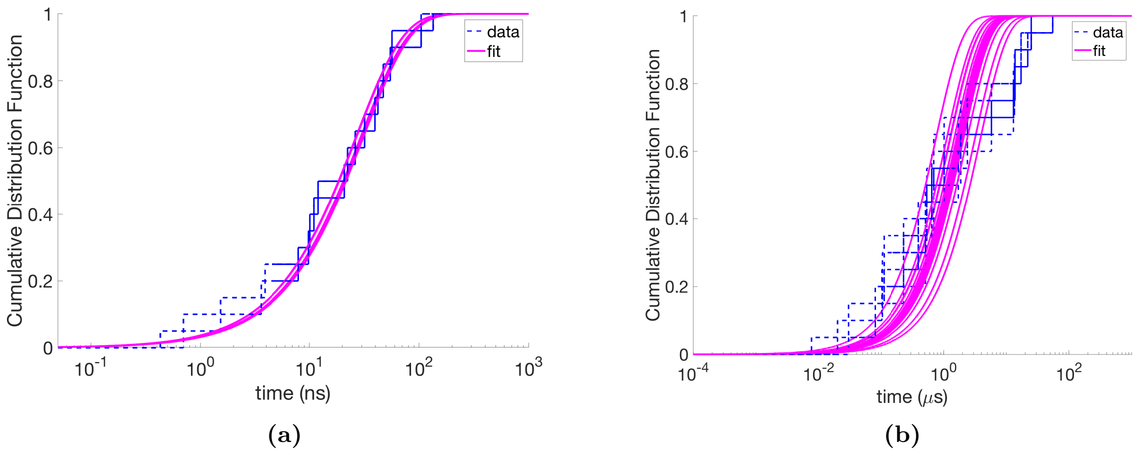


**FIG. 4:** Cumulative empirical probability distribution functions with their respective fits [2] for 20 sets of 20 transition times sampled with replacement from different independent dissociation events. Blue dashed line denote measured data, while magenta solid line denotes Poisson best-fit. (a) is for BUT drawing from total 21 independent observations, while (b) is for DAP drawing from total 25 independent observations.
